# Quantifying biofilm propagation on chemically modified surfaces

**DOI:** 10.1101/2020.11.20.391797

**Authors:** Michelle C. Halsted, Amber N. Bible, Jennifer L. Morrell-Falvey, Scott T. Retterer

**Affiliations:** The Bredesen Center, University of Tennessee, Knoxville TN, USA; Biosciences Division, Oak Ridge National Laboratory, Oak Ridge TN, USA; Center for Nanophase Materials Sciences, Oak Ridge TN, USA

## Abstract

Conditions affecting biofilm formation differ among bacterial species and this presents a challenge to studying biofilms in the lab. This work leverages functionalized silanes to control surface chemistry in the study of early biofilm propagation, quantified with a semi-automated image processing algorithm. These methods support the study of *Pantoea* sp. YR343, a gram-negative bacterium isolated from the poplar rhizosphere. We found that *Pantoea* sp. YR343 does not readily attach to hydrophilic surfaces but will form biofilms with a “honeycomb” morphology on hydrophobic surfaces. Our image processing algorithm described here was used to quantify this honeycomb morphology over time and displayed a logarithmic behavior in the propagation of the honeycomb biofilm. This methodology was repeated with a flagella-deficient *fliR* mutant of *Pantoea* sp. YR343 which resulted in reduced surface attachment. Quantifiable differences between *Pantoea* WT and Δ*fliR* biofilm morphologies were captured by the image processing algorithm, further demonstrating the insight gained from these methods.

## Introduction

Biofilm formation and disruption have tremendous implications across a breadth of fields, including human health, agriculture, and chemical processing, (Donlan, 2002; Bendaoud et al, 2011; Schultz et al. 2012; Petrova and Sauer, 2012; Kim et al., 2012; Wang et al., 2013; Fish, Osborn, and Boxall, 2017). Both physical and chemical cues mediate the processes by which these multicellular communities of bacteria attach and develop into complex architectures along natural and synthetic surfaces. Biofilm formation is often essential to the protection and propagation of microbial communities and can have significant consequences on the surrounding environment or host (Sutherland, 2001; Little et al., 2008; Hochbaum and Aizenberg, 2010; Berne et al., 2018). Developing strategies for selectively promoting or preventing biofilm formation requires a basic understanding of the factors that impact this process. Here we describe experimental methods that expand the current tools available for visualizing, analyzing, and quantifying biofilm formation and demonstrate the utility of these methods for analyzing biofilm formation in a microbial isolate from the poplar rhizosphere.

Physical and chemical factors collectively influence cell attachment and have a profound impact on biofilm propagation. Insight into these governing forces can be gained from fine control and manipulation of surface properties (e.g. hydrophobicity, surface roughness, surface topography) (Mozes et al., 1987; Costerton et al., 1999; Gottenbos et al., 2002; Garrett, Bhakoo, and Zhang, 2008; Glass et al., 2011; Epstein et al., 2011; Harimawan et al., 2011; Bendaoud et al., 2011; Friedladner et al., 2013; Tuson and Weibel, 2013). Many bacterial biofilms are difficult to visualize and interrogate in their natural environments, but bacteria can be cultured in nanofabricated and microfluidic platforms that offer exquisite control of surface properties through material deposition, definition of topography, and use of soft lithography (Friedlander et al., 2013; Hol and Dekker, 2014; Friedlander et al., 2015). When paired with appropriate microscopy and image processing, these platforms can facilitate visualization and quantitative descriptions of bacterial growth and biofilm formation.

Self-assembled monolayers are a promising approach to controlling surface properties for biofilm studies through the use of silane and thiol chemicals, tuned with different chain-end functional groups (Tan and Craighead, 2010; Glass et al., 2011; Privett et al., 2011; Wang et al., 2013; Tuson and Weibel, 2013; Friedlander et al., 2015; Maroni et al., 2015; Galbiati, 2016). The surface energy and reactivity can be readily modulated using commercially available reagents to modify silica, gold, and glass surfaces. One study utilized a hydrocarbon silane to survey attachment of *Staphylococcus epidermidis, Pseudomonas aeruginosa, Pseudomonas putida*, and *Escherichia coli* by performing cell counts from a video recording (Wang et al., 2013). Friedlander *et al*. utilized thiol self-assembled monolayers to evaluate the role of hydrophobicity on flagella adhesion with quartz crystal microbalance and dissipation (QCM-D), which leverages the extreme resonance sensitivity of a quartz crystal to quantify absorbed mass by change in resonance frequency (2015). While this is a suitable approach to quantify overall cell attachment and revealed the importance of *E. coli* flagella in cell attachment to hydrophobic surfaces, these methods do not describe biofilm morphology (Friedlander et al., 2015). Image processing can be applied to functionalized substrates to quantify cell attachment and morphology during the early stages of biofilm formation.

Imaging has been a hallmark of biological studies for centuries, providing qualitative descriptions of physical changes in biological systems since Hooke made the first observations of cells in the 17^th^ century. As more image processing tools have become readily available and more sophisticated, the extraction of quantitative information from these images can be used to explain intuitive trends with increased statistical rigor (Yang et al., 2001; Verma et al., 2012; Choudhry, 2016). Yang et al. demonstrated the ability of the Image Structure Analyzer software package to extract morphological characteristics of porosity and fractal dimension in monolayer biofilms (2001). Built-in particle analysis functions from ImageJ, MATLAB, or Python can perform automated cell counts, which can be further utilized to describe surface coverage, shape, and connectivity of developing films (Cai et al., 2011; Choudhry, 2016). We designed a semi-automated ImageJ script to quantify cell attachment and biofilm morphology from fluorescent microscopy images. When combined with automated image acquisition, these methods can quickly extract data from hundreds of images across multiple substrates. These data can then be used to describe cell attachment dynamics by different bacterial species across a variety of altered surface chemistries.

Our work examines the influence of hydrophilic and hydrophobic silane-treated substrates on *Pantoea* sp. YR343 attachment and biofilm propagation in static conditions. *Pantoea* sp. YR343 was isolated from the rhizosphere of *Populus deltoides* and has been shown to promote plant growth (Bible et al., 2016). Here, we describe propagation of biofilms with a honeycomb morphology produced by *Pantoea* sp. YR343 on hydrophobic surfaces, and leveraged ImageJ scripting to quantify this propagation mechanism. Flagella have been implicated in surface attachment, thus we examined a flagella-defective mutant of *Pantoea* sp. YR343 to quantify the impact of flagella on biofilm propagation with these methods (Friedlander et al., 2015). Compared to the wildtype strain, the flagellar mutant showed delayed and reduced cell attachment and differences in biofilm morphology, consistent with consequences of motility and surface adhesion from a compromised flagellum.

## Results & Discussion

*Pantoea* sp. YR343 is a rod-shaped, flagellated, gram-negative bacterium which was isolated from the Poplar rhizosphere and has been shown to exhibit biofilm formation, swimming motility, and surface motility (Bible et al., 2016). *Pantoea* sp. YR343 has been engineered to express green fluorescent protein (GFP), facilitating the use of fluorescence microscopy for quantifying cell attachment and surface coverage. Silicon dioxide coated substrates were modified to examine the impact on cell attachment using the following chemistries: a fluorinated chain, a hydrocarbon chain, an ester, and an amine group. These substrates were characterized by hydrophobicity, as indicated by the water contact angle measurement in Table 1. Materials with a contact angle greater than 90° are considered hydrophobic.

**Table 1:** Summary of silane acronyms and water contact angel measurements.

|  | PFOTS | OTS | MTMS | APTMS |
| --- | --- | --- | --- | --- |
| **Chemical Name** | Trichloro (1H, 1H, 2H, 2H-perfluorooctyl) silane | n-octadecyl (trimethoxy) silane | Methoxytriethylenoxypropyl- trimethoxy silane | 3-aminopropyl trimethoxy silane |
| **Surface Property** | Hydrophobic | Hydrophobic | Hydrophilic | Hydrophilic |
| **Water Contact Angel** | 104° | 93° | 45° | 53° |

*Pantoea* sp. YR343 attachment to functionalized substrates was tested under static conditions. Substrates were submerged in 3 mL of R2A culture medium inoculated with *Pantoea* sp. YR343-GFP, at an optical density of 0.1 (at 600 nm). Substrates were removed from the culture at selected time points and rinsed with 10 mL distilled water to remove loosely attached cells. Care was taken to apply the water near the edge of the substrate and flow water across the sample. The substrate was dried with pressurized air to minimize drying artifacts and subsequently imaged with a fluorescence microscope (Figure S.1).

Fluorescence images of the early biofilm were collected from multiple positions across each sample to capture representative cell behavior. An ImageJ script applied a built-in threshold function, Huang or Default, to generate a binary image (Figure 2). The binary image produced from each threshold function was evaluated, selected, and manually adjusted to most accurately reflect surface area coverage, i.e. cell attachment (Choudhry, 2016). A binary function inverted the image to facilitate counting and sizing of the gaps present in the honeycomb morphology (Figure 2). Biofilm morphology could not be computed directly because of the interconnected honeycomb pattern. The ImageJ particle analysis function identified the (black) gaps in the inverted image as the object of interest (Figure 2). This “gap analysis” quantified the gap size, number of gaps, and surface area coverage (%). Subtraction of the gap area coverage from 100 yielded the percentage of surface area covered by the cells. For context, *Pantoea* sp. YR343 is 1-2 μm in length, approximately 0.5 μm in width, and 18-38 pixels in size, thus the gap analysis limit was set to 18 pixels (this analysis focuses on the morphology features with sizes greater than that of a single cell); the image is approximately 380,000 pixels.

**Figure 2:**
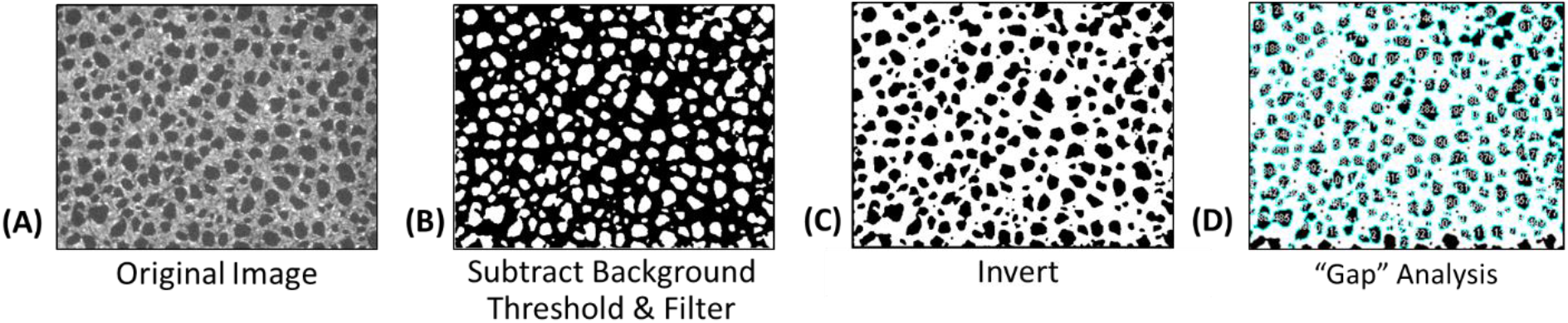
Image processing and quantification of honeycomb biofilm pattern. Cells are grey in the original image (A) and black in the binary image (B). The image is inverted (C), and the particle analysis function is applied to gaps (black) in the image (D). *Pantoea* sp. YR343 is 1-2 μm in length, approximately 0.5 μm in width, which equates to 18-38 pixels. The lower limit of the particle analysis function is set to 18 pixels, the smallest measure of a single *Pantoea* sp. YR343.

Figure 3 summarizes the effect of hydrophobicity on cell attachment. After twenty hours, *Pantoea* sp. YR343 biofilm surpassed 70% area coverage on the hydrophobic surfaces (PFOTS and OTS), with neglible attachment to the hydrophillic surfaces (APTMS and MTMS). There was also neglible attachment to the control surfaces of glass, silicon, and quartz (data not shown). These results are consistent with the literature as many bacterial species have been shown to be negatively charged and favor attachment to hydrophobic, neutral surfaces (Donlan, 2002; Mai and Corner, 2007; Song, Koo and Ren, 2015; Berne et al., 2018).

**Figure 3:**
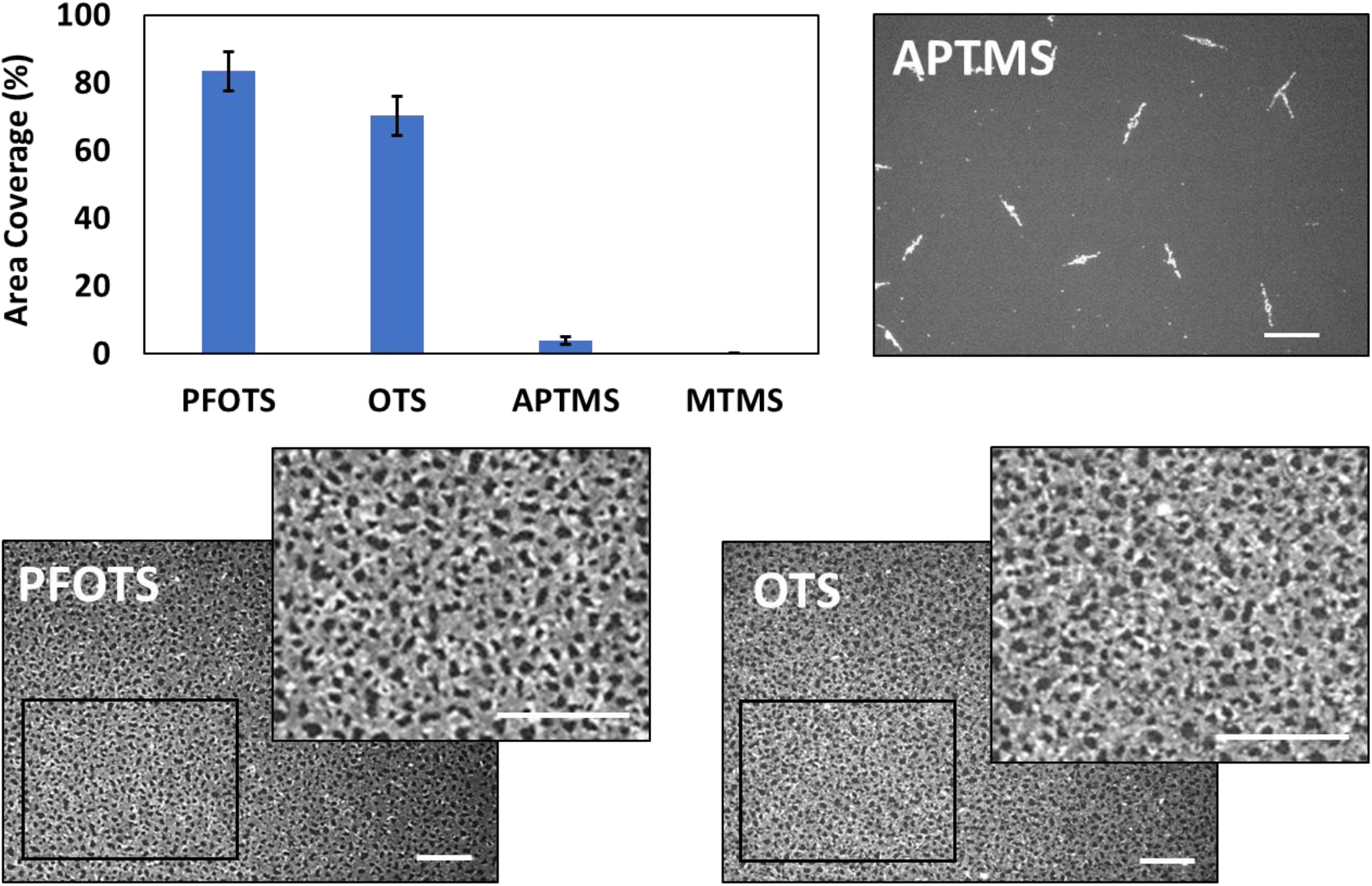
*Pantoea sp*. YR343 area coverage on hydrophobic and hydrophilic surfaces after 20 hour: Pantoea sp. YR343 attachment to Trichloro(1H,1H,2H,2H-perfluorooctyl) silane (PFOTS), n-octadecyl (trimethoxy) silane (OTS), 3-aminopropyl trimethoxy silane (APTMS), and Methoxytriethyleneoxypropyl-trimethoxy silane (MTMS). Scale bar 25 μm.

### Propagation of Honeycomb Biofilm Morphology

Honeycomb biofilm morophology (also referred to as web-like, net-like, networks, and branching morphology) is not unique to *Pantoea* sp. YR343, and has been previously observed in the literature, under a variety of conditions (e.g. static conditions, fluid flow; stainless steel coupon, polystyrene microtiter plate; mixed biofilm, pure culture; wet biofilm, dried biofilm), with both Gram-negative and Gram-positive bacterial strains (Donlan et al., 2002; Marsh, Luo and Wang, 2003; Takhistov and George, 2005; Bridier et al., 2010; Serra et al., 2013; Mosquera-Fernández, 2014; Guilbaud et al., 2015; Chavant et al., 2002). Custom image processing scripts quantified the evolution of *Pantoea* sp. YR343 biofilm morphology based on the number and size of gaps observed in the honeycomb pattern. Figure 4 depicts *Pantoea* sp. YR343 attachment to PFOTS with an average area coverage for each time point, with representative images to illustrate the evolution of the honeycomb biofilm morphology.

**Figure 4:**
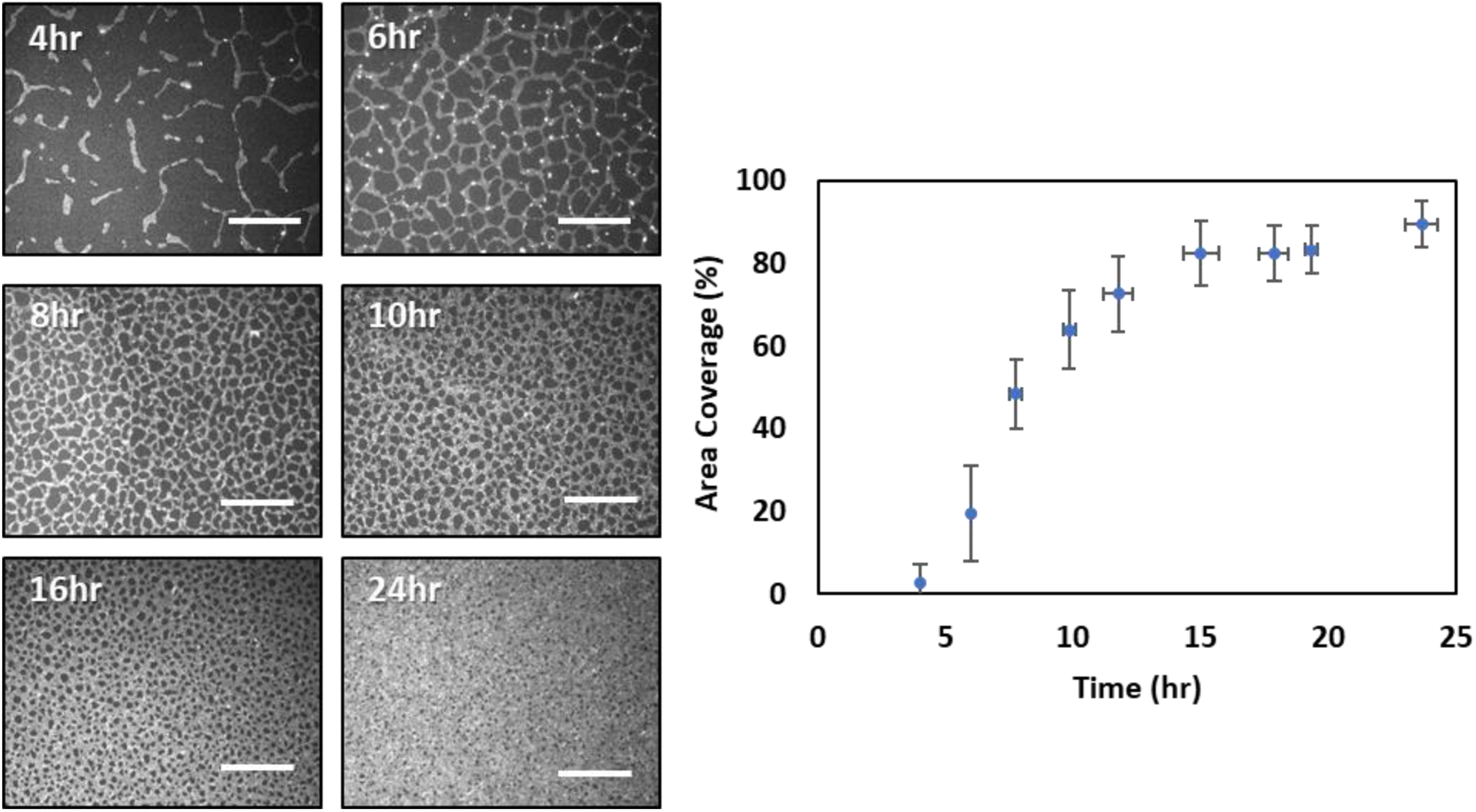
*Pantoea sp*. YR343 attachment to Trichloro(1H,1H,2H,2H-perfluorooctyl) silane (PFOTS), average area coverage for each time point. Image scale bar 50 μm, error bar: 1 standard deviation area coverage (%).

As shown in Figure 4, the *Pantoea* sp. YR343 biofilm begins with linear branches of cells which extend in length and intersect with other branches to create a net-like or honeycomb appearance. The honeycomb gaps are segmented with branches of cells as the biofilm continues to propagate. This consequently decreases the gap size and increases the number of gaps in the honeycomb biofilm. The gaps become smaller and many of the gaps are eventually filled by cells. Substaintially fewer gaps remain in the 24-hour dataset. Gaps less than the cell size may be present, but have been excluded from these analyses due to user-defined minimum allowable gap sizes. These visual observations are supported with quantitative information extracted from the image via gap analysis (Figure 5).

**Figure 5:**
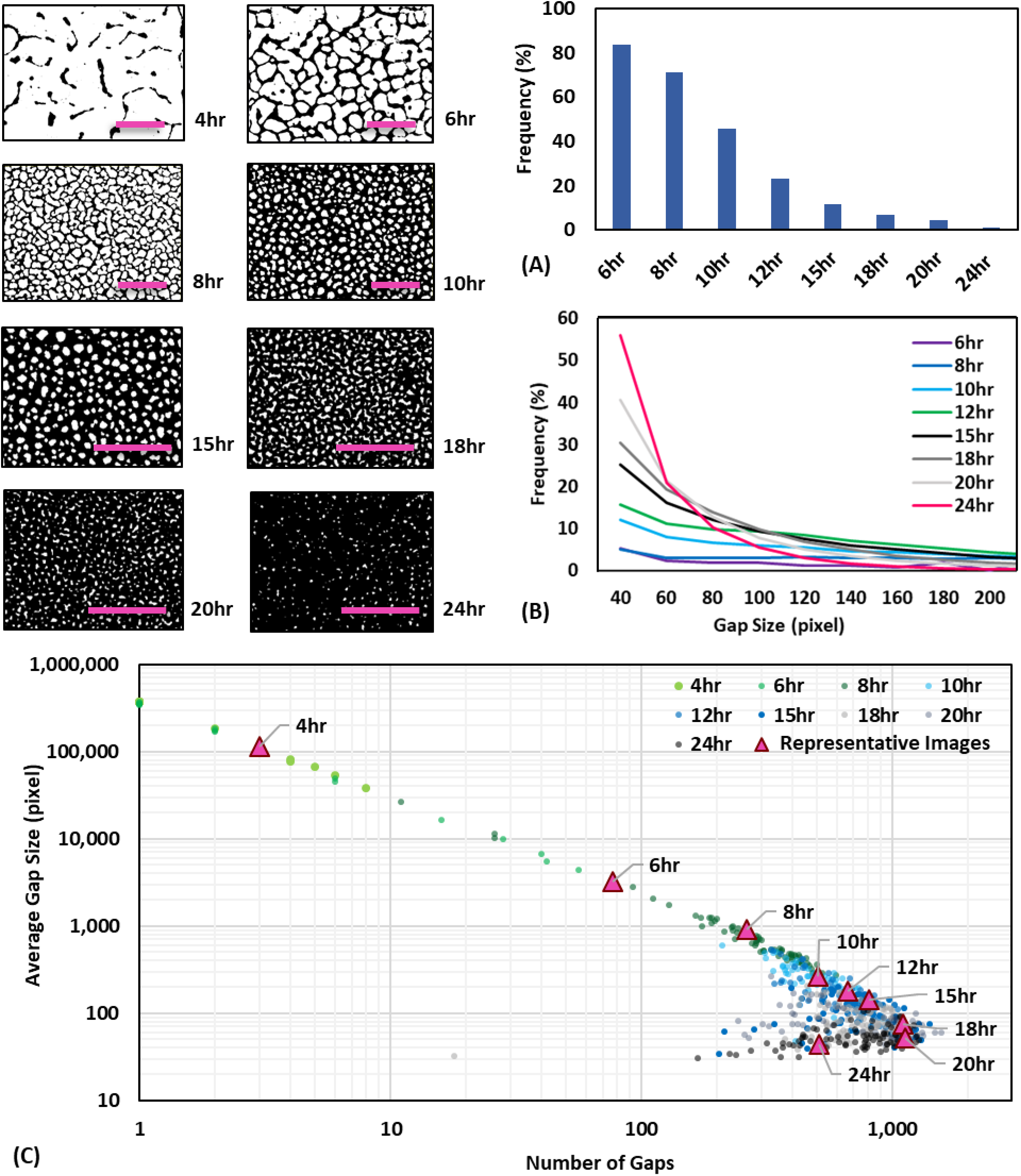
Characterization of *Pantoea* sp. YR343 morphology with gap size and number. A) Distribution of gap size across *Pantoea* sp. YR343 dataset (percentage of gaps exceeding 200 pixels size is not shown in plot); 45 pixels corresponds to 1 **μ**m. B) Inset depicts the measure of gap sizes in pixels. C) Percentage of gap sizes greater than 200 pixels for each time point. D) Relationship between average gap size and number per image, represented by a data point. Representative images from each time point correspond to the pink triangle data point on the plot, scale bar is 50μm.

Figure 5, A shows the percentage of gaps greater than 200 pixels for at each time point. Figure 5, B shows the gap size distribution at each time point, based on a range of 18-200 pixels. The 4-hour dataset is not included in the analyses because the morphology is dominated by unconnected, linear branches of cells at this time point. The few gaps that formed during the 4-hour time point are on the order of 10,000 and 100,000 pixels.

The percentage of *Pantoea* sp. YR343 gaps incrementally decrease as the gap size bin increases, and this is consistent with the logarithmic relationship between average gap size in an image and the number of gaps across time points (Figure 5, A, C). Approximately 40% of the 20-hour dataset is comprised of gaps with less than 40 pixels in size, and this jumps to 60% in the 24-hour dataset. The high percentage of small gap sizes fits with 90% surface area coverage after 24 hours (Figure 4). While these images are of a monolayer *Pantoea* sp. YR343 biofilm, confirmed by Scanning Electron Microscopy in Figure (SEM), it is highly plausible that the rinse step removed loosely attached cells from a multi-dimensional biofilm matrix (Supplementary Materials Figure S.2). In a three-dimensional biofilm, these honeycomb gaps would result in a porous biofilm, facilitating mass transfer of nutrients, waste, and oxygen (Donlan, 2002; Flemming and Wingender, 2010; Petrova and Sauer, 2012; Mosquera-Fernández et al., 2014).

Figure 5, D illustrates an exponential decrease in average gap size accompanied by an exponential increase in the number of gaps. From this logarithmic behavior, we infer *Pantoea* sp. YR343 cells segment gaps in the honeycomb biofilm as part of the biofilm propagation mechanism where one large gap becomes two small gaps, and the average size of these two gaps equates to half the size of the large gap. This behavior follows a slope of −1, which is the approximate slope of the *Pantoea* sp. YR343 dataset in Figure 5, D between 4 and 15 hours. Many, but not all, of the gaps in the honeycomb biofilm eventually become so small that they are filled by cells (18-24 hours), and this consequently decreases the number of gaps (i.e. creates a bend in the dataset). The representative images show in Figure 5, D correspond to the triangle data points and offer a simplified example. Figure S.3 offers a complementary plot relating average gap size for each dataset with respect to time.

### Propagation of Flagella Mutant Biofilms

Flagella play a key role in the initial stages of biofilm formation. In addition to providing a motor for swimming and surface motility, flagella can mediate attachment by overcoming repulsive forces near the surface (Lemon et al., 2007; Petrova and Sauer, 2012; Friedlander et al., 2015; Kearns, 2010; Guttenplan and Kearns, 2013; Berne et al., 2018). Flagella increase the surface area of attachment and have been shown to anchor cells to surfaces (Lemon et al., 2007; Tuson and Weibel, 2013; Friedlander et al., 2015; Berne et al., 2018). Like *Pantoea* sp. YR343, *Listeria monocytogenes* form honeycomb biofilms and flagella have been demonstrated to play an integral role in this morphology as the absence of flagella resulted in unstructured biofilms (Gauilbaud et al., 2015). To test whether our method could distinguish different biofilm morphologies, we examined biofilm formation using a *Pantoea* sp. YR343 mutant defective in flagella assembly due to deletion of the gene encoding FliR (Figure 6).

**Figure 6:**
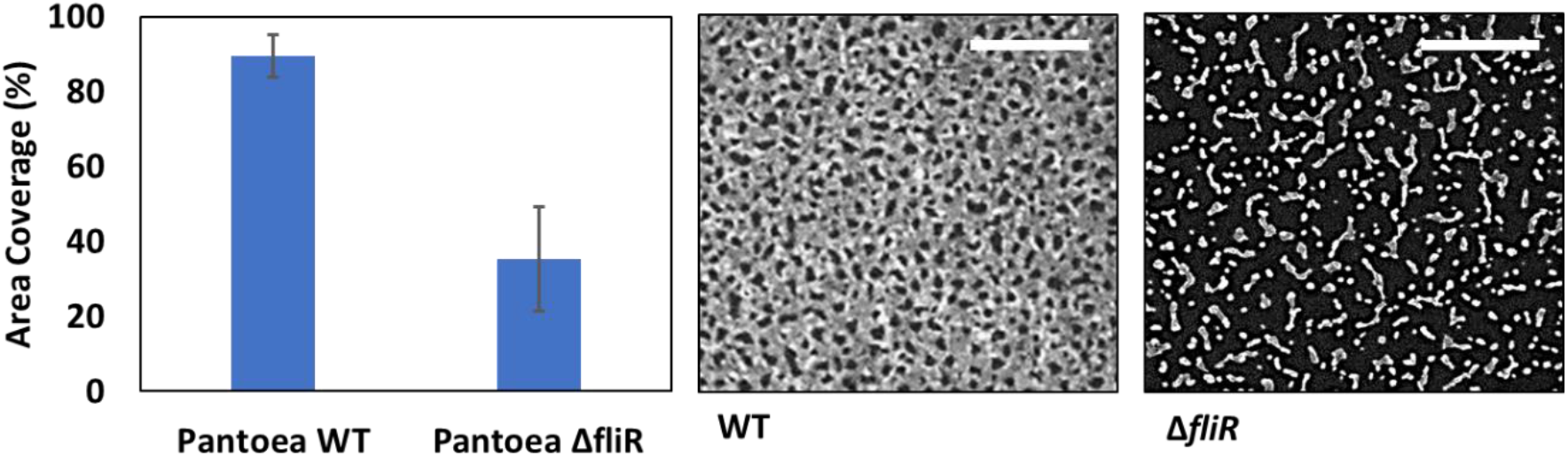
Effect of *Pantoea* sp. YR343 flagella on attachment to PFOTS, scale bar 25 μm. Error bar: 1 standard deviation.

FliR is a conserved integral membrane protein in the basal body complex that plays a key role in the structure and function of the flagellar export apparatus (Nakamura and Minamino, 2019). Using this mutant, we found that the area coverage of Δ*fliR* biofilms is approximately half that of wildtype biofilms, yet the standard deviation is approximately doubled (Figure 6). When examined with SEM, no flagella were observed on the mutant cells (Supplementary Materials, Figure S.4, S.5). The variation may be explained by the decrease in surface adhesion due to loss of the flagella adhesin. When examined with scanning electron microscopy, no flagella were observed on the mutant cells (Supplementary Materials, Figure S.3, S.4). Consistent with previous reports, a lack of flagella or defects to the flagella adhesin likely explains the Δ*fliR* area coverage and variation (Lemon et al., 2007; Serra et al., 2013; Friedlander et al., 2013; Friedlander et al., 2015). Relatively large sections of the biofilm appeared to detach during the rinse step, and these experimental observations align with the notion that the Δ*fliR* biofilm lacks sufficient adhesion.

Using our image processing algorithm, we quantified *Pantoea* sp. YR343 Δ*fliR* biofilm morphology compared to wild type cells (Figure 7). This analysis shows that there are dramatic differences in the gap size distribution, with many of the gaps in the *Pantoea* sp. YR343 Δ*fliR* dataset exceed 200 pixels (Figure S.6, A). Interestingly, gap sizes below 200 pixels in the Δ*fliR* dataset are evenly distributed across time points (Figure S.6, B). This is consistent with Figure 7, A which shows a scattered relationship between average gap size and number that does not change with respect to time. In other words, the *Pantoea* sp. YR343 Δ*fliR* dataset may follow the same spatial trend as *Pantoea* sp. YR343 WT but does not follow the temporal trend (Figure 7, A).

**Figure 7:**
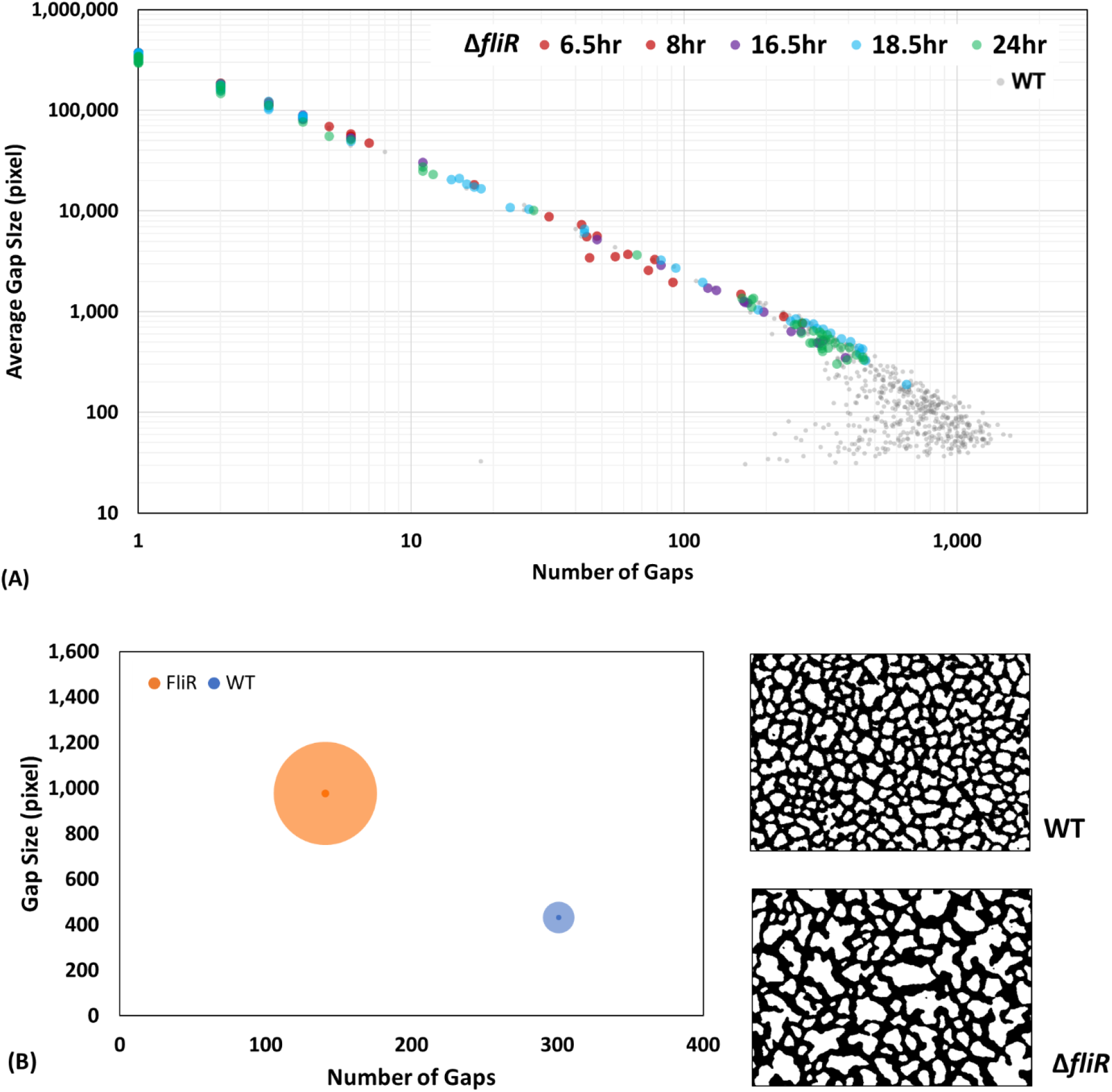
Differences between *Pantoea* sp. YR343 WT and *Pantoea* sp. YR343 Δ*fliR* biofilm morphology. A) Relationship between **Δ***fliR* average gap size and number per image, represented by each data point and overlaid on WT dataset. B) Quantitative comparison of biofilm morphology where bubble size denotes standard deviation in gap size, with representative images for *Pantoea* sp. YR343 WT (53% area coverage) and *Pantoea* sp. YR343 Δ*fliR* (52% area coverage).

Figure 7, B captures the differences in the *Pantoea* sp. YR343 Δ*fliR* and WT morphology by comparing metrics of two (representative) biofilm images with equal surface area coverage. The radius of the bubble corresponds to the standard deviation in the image gap size, and the dot in the center of the bubble indicates the average gap size for each image. The radius of the bubble is independent of the gap number and is represented by bubble position.

Confocal microscopy has the benefit of capturing three-dimensional data and electron microscopy can interrogate biofilms on the sub-micron scales (Figure 8). The honeycomb biofilm is visible in each of these microscopy images, and the three-dimensional profile of the biofilm appears to have a dense layer of cells at 24 hours. We believe this profile consists of loosely attached cells, possibly accrued from settling artifacts, and these cells were washed away during the rinse step. We also found that temperature affects the rate of biofilm propagation, and future work be expanded to inclcude temperature and the three-dimensional biofilm studies.

**Figure 8:**
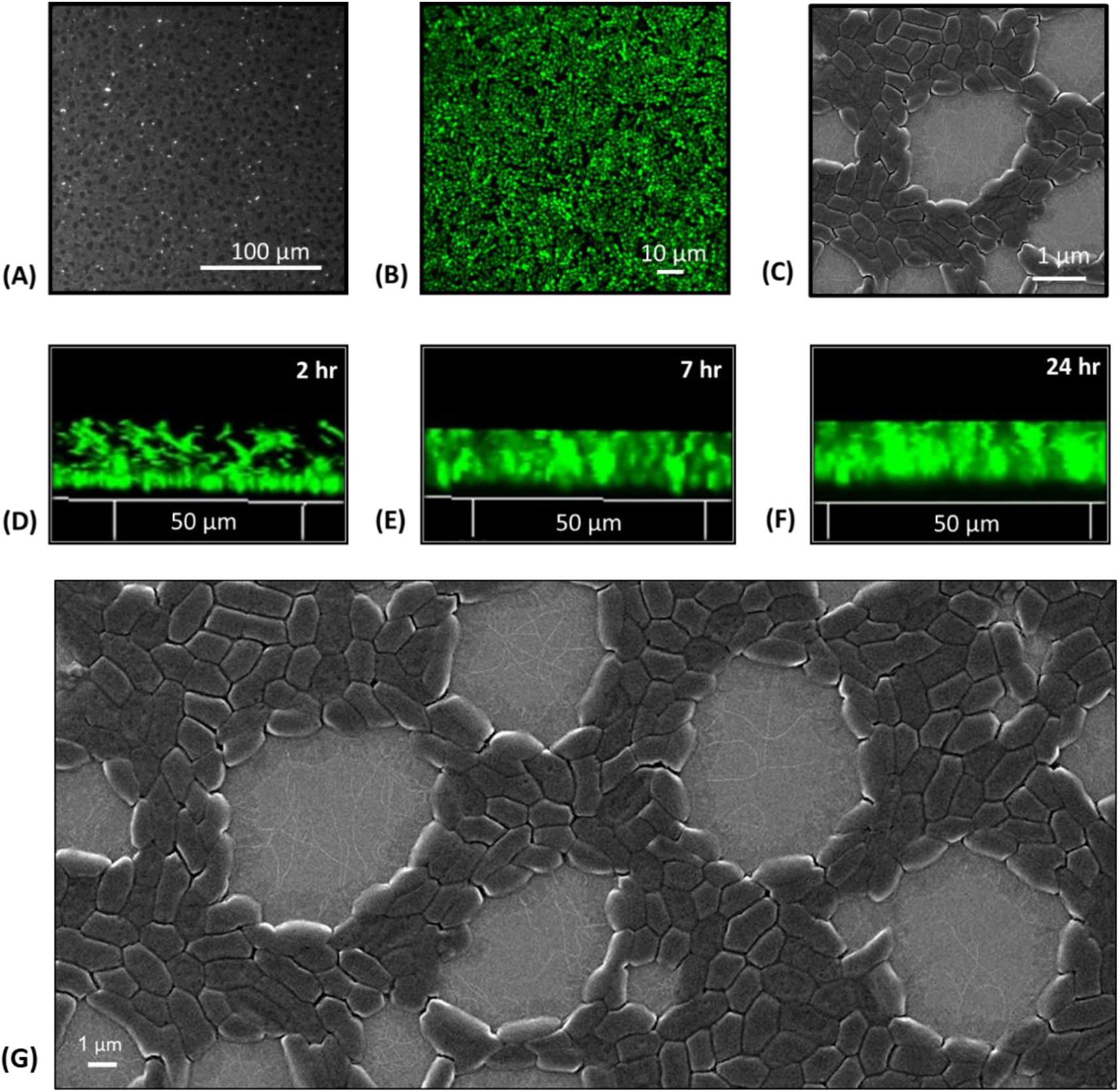
An evaluation of *Pantoea* sp. YR343 biofilm propagation on PFOTS-silicon substrate using different microscopy methods. A) Fluorescence microscopy after 10 hours attachment. Sample was rinsed with 10 mL DI water and dried with pressurized air. B) Scanning Electron Microscopy after 10 hours of attachment. Sample was rinsed with 10 mL DI water, dried with pressurized air, and coated with 5 nm gold. C) Wet biofilm at 13 hours, Confocal Laser Scanning Microscopy. D) Vertical profile of *Pantoea* sp. YR343 biofilm at 2 hours. E) Vertical profile of *Pantoea* sp. YR343 biofilm at 7 hours. F) Vertical profile of Pantoea sp. YR343 biofilm at 24 hours. G) Pantoea sp. YR343 biofilm propagation on PFOTS-Si substrate, 24 hours (Zeiss Scanning Electron Microscope, 5 nm gold coating).

Scanning electron microscopy images revealed a monolayer of cells in the honeycomb biofilm with flagella spanning the gaps (Figure 8, G). Flagella often appeared to be intertwined, and occasionally chains of cells extended from one side of the biofilm, into part of the gap. These cells appeared to be in the process of segmenting the gap when the biofilm was removed from the culture. The presence of the flagella in the middle of the biofilm gap suggests the flagella serve as an attachment point for motile cells, and this could explain why the flagella defective mutant biofilms propagate on a much slower timeline and with seemingly lower adhesion. SEM images show the flagella spanning across the biofilm gaps, thus its feasible that the flagella guide the nascent cells to segment the gap by providing additional points of attachment (Figure 8, G).

## Conclusions

Our methods leverage the ease of fluorescence microscopy to generate large datasets, across multiple surfaces and samples, necessary for capturing the natural variation and distribution of biological phenomenon. This approach will complement the qualitative data gathered from confocal and scanning electron microscopy, which are resource intensive and time consuming. We demonstrated how this alternative biofilm assay quantifies early biofilm formation on silane-treated surfaces. The semi-automated image processing script captures numerical data in biofilm morphology and unveiled considerable differences in the morphology of *Pantoea* sp. YR343 WT and *Pantoea* sp. YR343 Δ*fliR* biofilms. Additionally, our approach quantifies spatial temporal data qualitatively observed in microscopy images of biofilm propagation. Numerical data enables statistics and regression analysis, which can in turn be incorporated into computer simulations. Collectively, novel and traditional methods will be essential in unraveling the complexities of biofilm formation.

## Methods

### Surface chemistry modification

Silicon wafers with silicon dioxide coating (Silicon Quest), were diced into 20 mm by 20 mm square chips. The chips were cleaned with pressurized air with a 0.2 μm filter, followed by a minimum of 5 minutes in a Harrick Plasma PDC-001 air plasma cleaner (Ithaca, NY). Vapor deposition was performed in an enclosed, glass dish on a hot plate with the following methods: 20μL per 80 cm^2^ trichloro(1H,1H,2H,2H-perfluorooctyl) silane (PFOTS) (Sigma-Aldrich, St. Louis, MO) for 4 hours at 85°C; 40μL per 80 cm^2^ 3-aminopropyl trimethoxysilane (APTMS) (Gelest, Morrisville, PA) for 2 hours at 150°C; 40μL per 80 cm^2^ n-octadecyl (trimethoxy)silane (OTS) (Gelest, Morrisville, PA) for 2 hours at 150°C, followed by 2 hours no heat; 4 hours 65°C Methoxytriethyleneoxypropyl-trimethoxysilane (MTMS) (Gelest, Morrisville, PA), followed by 1 hour at 115°C.

### Bacterial Culture and Device Testing

Engineered strains of *Pantoea* sp. YR343 expressing green fluorescent protein were engineered by expression of EGFP from a Gate-way modified pBBR1-MCS5 plasmid, maintained with 10 μg gentamycin, ml (Bible et al., 2016). *Pantoea* sp. YR343 were inoculated in R2A liquid medium (from a plate of R2A agar) and grown to stationary phase overnight. A 1:100 dilution was performed, and the culture was grown to early exponential phase (approx. 4hrs) and diluted to an optical density (OD) of 0.1, verified with a BioTek Synergy 2 microplate reader, 600 nm.

The silane-treated substrates were each placed in concave dishes and filled with 3 mL of Pantoea sp. YR343 liquid culture, 0.1 OD. Upon inoculation, the dishes were covered and incubated at for a specified amount of time. Tweezers were used to remove the substrate from the liquid culture at a designated time point. The substrate was rinsed with 10 mL DI water to remove loosely attached cells, and dried with pressurized air, 0.2 μL filter, to minimize drying artifacts. These experiments were conducted at a room termperature around 60 °C.

### Sample Imaging

Image data was collected with an Olympus IX51 microscope (Shinjuku, Tokyo) complete with epifluorescence using a Chroma 41001FITC (Bellows Falls, VT) filter cube (480nm excitation band pass filter with a 40nm band width and 535nm emission band pass filter with a 50nm band width). FEI Novalab 600 Dual-Beam System was used to collect Scanning Electron Microscopy (SEM) images of the *Pantoea* sp. YR343 cell attachment. Confocal fluorescence microscopy was performed using a Zeiss LSM710 confocal laser scanning microscope with a Plan-Apochromat 63x/1.40 oil immersion objective (Carl Zeiss Microimaging, Thornwood, NY).

### Image Processing and Cell Quantification

Images were analyzed in Fiji ImageJ. Image processing was executed with semi-automated scripts to generate binary images. Scripts were tailored for each stack of images, but all scripts applied built-in functions for background subtraction, threshold, and filters. The binary images were manually adjusted to match the original image. To extract numerical data from the honeycomb biofilm, another image script inverted each image and applied the built-in particle analysis function with a limit of 18 pixels. By inverting the images, the biofilm gaps became the region of interest and facilitated collection of morphology data. The built-in particle analysis function collected data on total particle area, percent area coverage, average particle size, and the number of particles. Cell area coverage was calculated by subtracting the gap area coverage percent from one hundred.

### Mutant strain construction

These methods are described fully in Bible et al. (2020). In short, biparental mating introduced the plasmid pRL27 and encoded a mini-Tn5 transposon into *Pantoea* sp. YR343 (DGC2884 pSRK-Gm). *Pantoea* sp. YR343 was grown in presence of kanamycin (50 μg mL^−1^) and gentamycin (10 μg mL^−1^) to remove of *E. coli* strain EA145 (Bible et al., 2020). The transposon library was screened, and genomic DNA was isolated from each mutant using the Promega Wizard Genomic DNA Extraction Kit, (Bible et al., 2020). Colonies were picked, the plasmid DNA was isolated with the QIAprep Spin Miniprep Kit (Qiagen), and plasmids were sequenced at the Molecular Biology Resource Facility at the University of Tennessee, Knoxville (Bible et al., 2020). The plasmids were sequenced using the primers tpnRL17-1 and tpnRL13-1, and the results were analyzed using BlastX from NCBI to identify the region of DNA flanking each transposon (Bible et al., 2020).

## Acknowledgements

This work was supported by the Ofice of Science, Biological and Environmental Research, as part of the Plant Microbe Interfaces Scientific Focus Area (http://pmi.orn.gov). Scanning electron microscopy was carried out at the Center for Nanophase Materials Sciences, which is a DOE Office of Science User Facility.

