## Supplementary Materials for "Quantifying biofilm propagation on chemically modified surfaces"

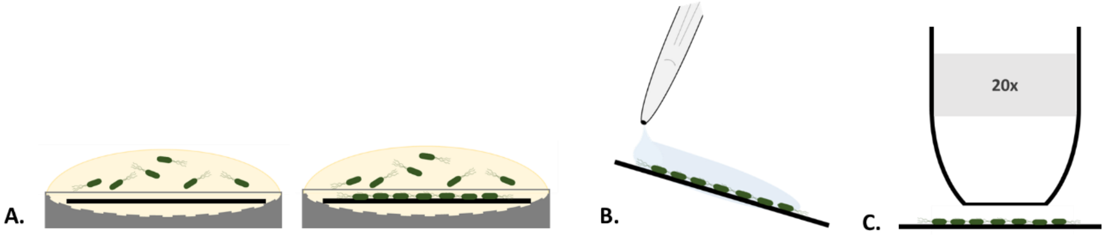


**Figure S.1: Biofilm assay methods. A) Substrates were submerged in 3 mL of R2A growth media inoculated with *Pantoea* sp. YR343 at an optical density ($OD_{600}$) reading of 0.1, and substrates were incubated under static conditions to allow surface attachment and biofilm formation; B) Substrates were removed at selected times and rinsed with 10 mL of DI water, and dried with pressurized air (0.2 µm filter); C) Imaging was carried out using a with 20x objective.**

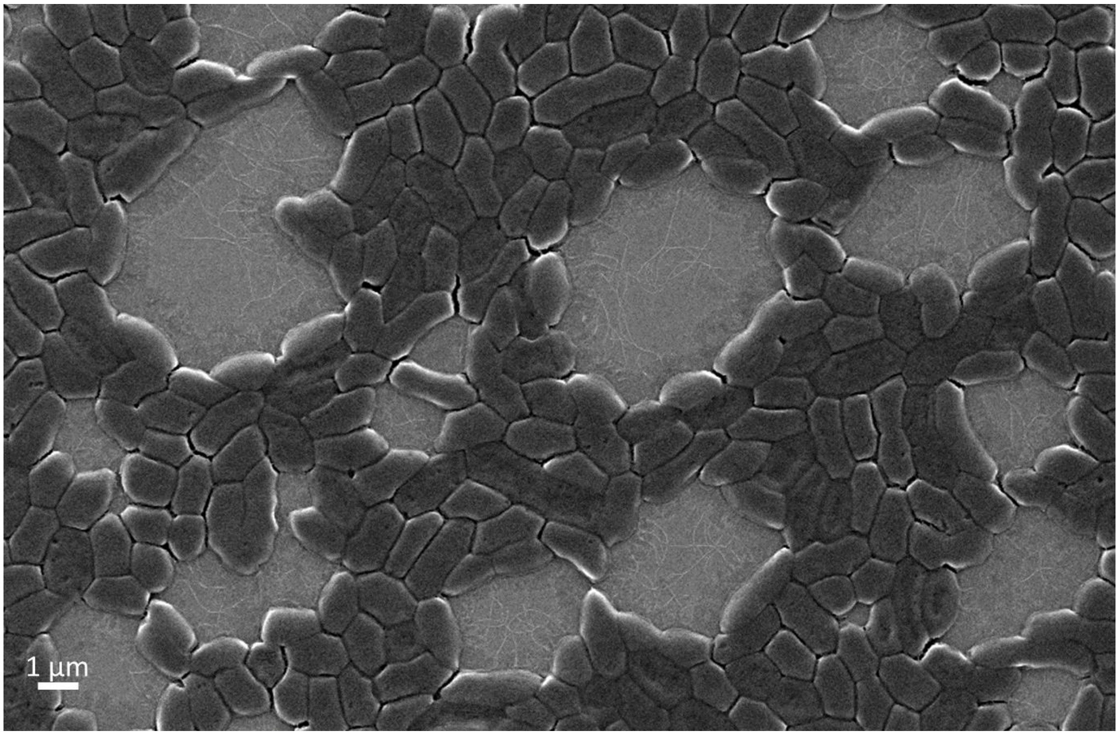


**Figure S.2: *Pantoea* sp. YR343 biofilm propagation on PFOTS-silicon substrate after 10 hours (Scanning Electron Microscopy, 5 nm gold).**

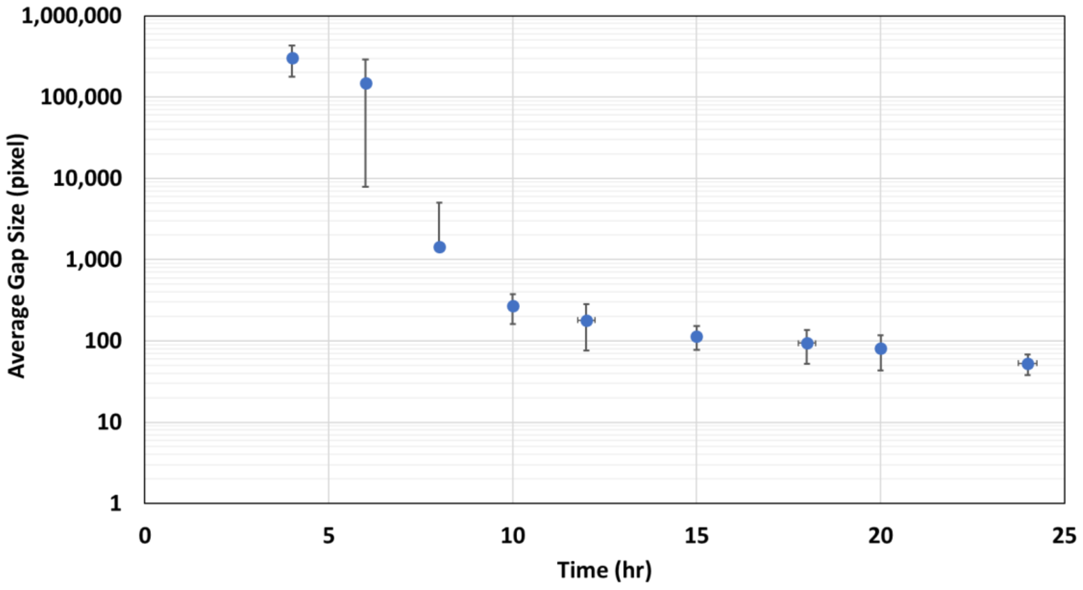


**Figure S.3: Average gap size for each dataset with respect to time.  Error Bar: 1 Std Dev.**

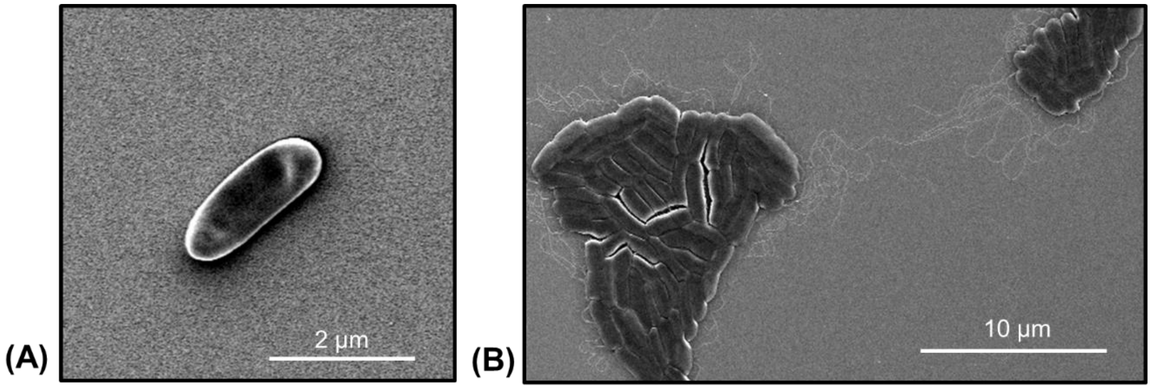


**Figure S.4: Early attachment of *Pantoea* sp. YR343 WT and flagella mutants to PFOTS-Si substrate (Zeiss Scanning Electron Microscope, 5 nm gold coating); A) *Pantoea* sp. YR343 Δ*fliR*, 3 hours. B) *Pantoea* sp. YR343 WT 4 hours.**

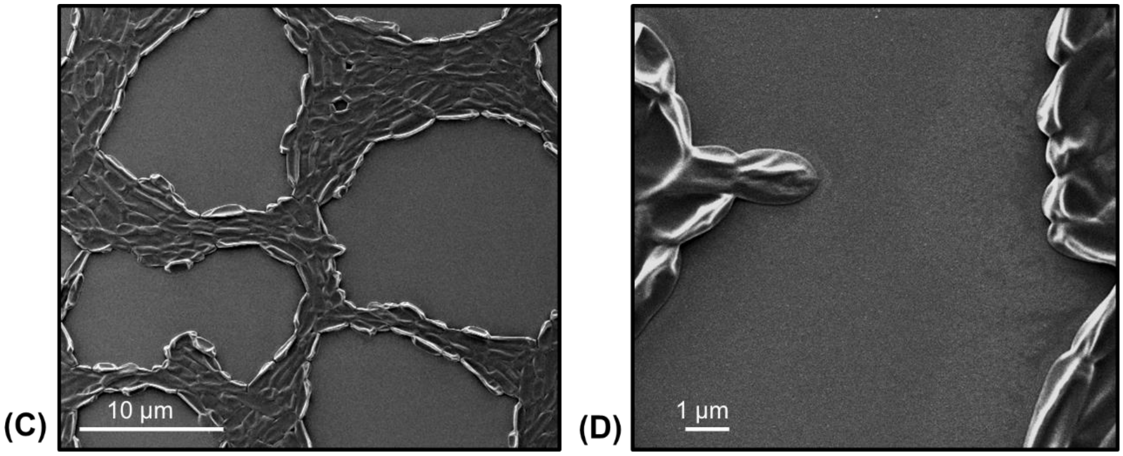


**Figure S.5:** ***Pantoea*** **sp. YR343** ***ΔfliR*** **mutant on PFOTS-Si substrate, 24 hours (Zeiss Scanning Electron Microscope, 5 nm gold coating).**

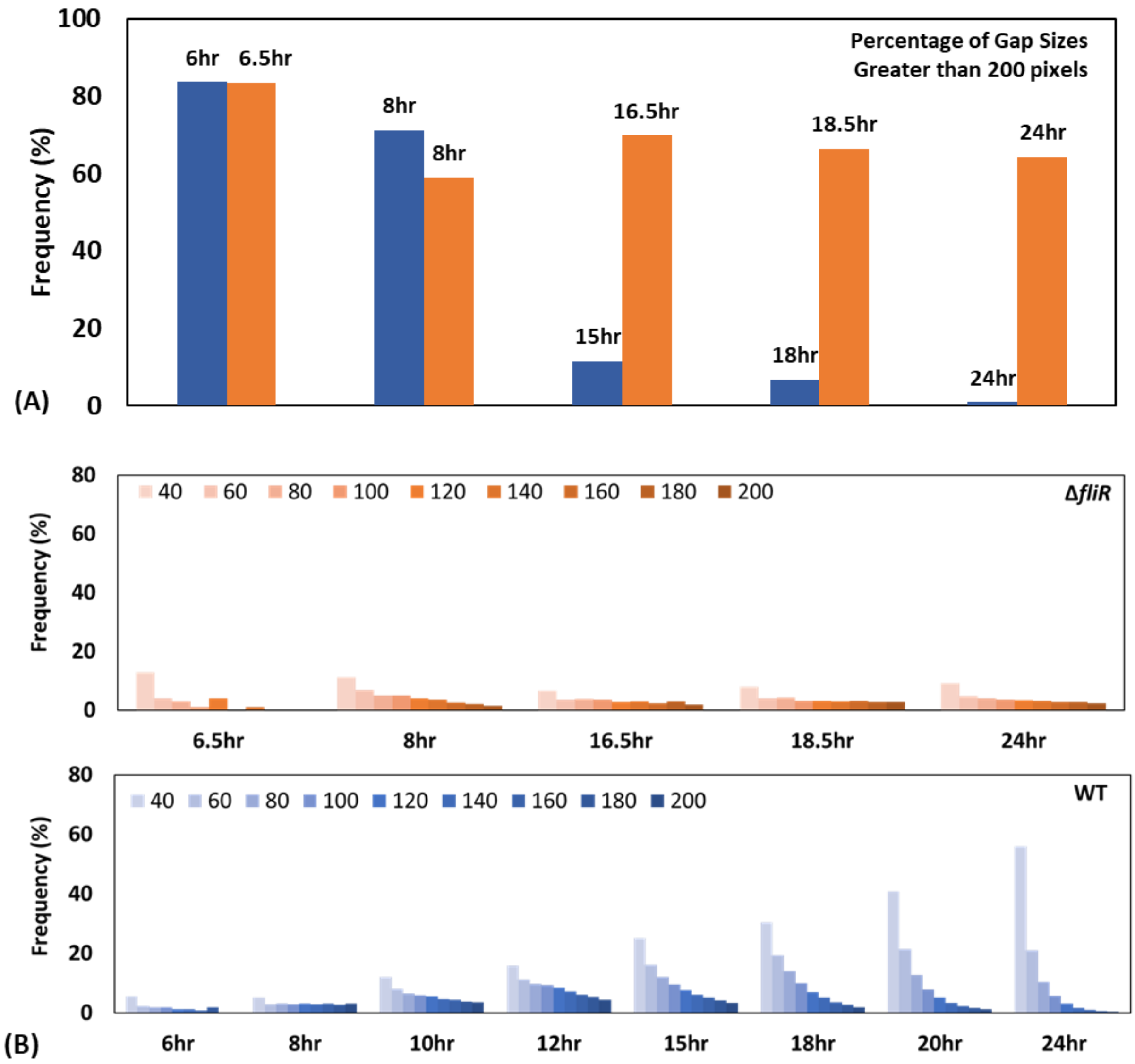


**Figure S.6: A) Comparison of WT (blue) and Δ*fliR* (orange) datasets, percentage of gap sizes greater than 200 pixels. B) Distribution of *Pantoea* sp. YR343 Δ*fliR* (top) vs. *Pantoea* sp. YR343 WT (bottom). Percentage of gaps exceeding 200-pixel size is not shown.**
